# Neuronal synapses are stabilised by activity-dependent vesicular release from oligodendrocyte precursor cells

**DOI:** 10.64898/2026.02.03.701391

**Authors:** Emma Dumble, Denis Yuan, Eneritz Agirre, Patricia Bispo, Laura Hoodless, Yan Xiao, Gonçalo Castelo-Branco, Tim Czopka

## Abstract

Activity-dependent regulation of synapses has long been attributed to interactions between neurons alone. Here, using *in vivo* imaging in the zebrafish visual system we show that one type of glia called the oligodendrocyte precursor cell (OPC) closely interacts with pre-synapses of retinal ganglion cell (RGC) axons, and exhibits activity-dependent Vamp2/3-mediated vesicular release. Critically, blocking OPC vesicle release using botulinum toxin, and silencing OPC activity integration through dampening calcium signalling, destabilises RGC pre-synapses. These observations suggest that OPCs provide a non-neuronal pathway for activity-dependent feedback to shape circuit wiring.

## Main text

It is established that the fine-tuning of neuronal connectivity is governed by activity-dependent feedback mechanisms, which can structurally manifest in the selective formation, stabilisation and elimination of synapses to strengthen or weaken specific connections of a circuit ^1,2^. In recent years glial cells are increasingly implicated in contributing to the regulation of circuit connectivity ^3–8^, challenging the traditional view that the tuning of synaptic connectivity is reliant on communication between neurons alone. Here, we explored if one specific type of glia called the oligodendrocyte precursor cell (OPC) may directly feedback to neurons in an activity dependent manner to tune synapse stability.

OPCs persist throughout the CNS from development through adulthood and ageing where they maintain elaborate process networks that tile the tissue ^9^, and it was recently shown that they have mature, myelination-independent functions in regulating neuronal connectivity ^3–6^. Moreover, they express a wide range of neurotransmitter receptors and voltage gated ion channels, and even receive direct post-synaptic contacts from neurons ^10–12^. Together, this gives OPCs the principal capacity to participate in activity-dependent circuit tuning.

We have previously established that the optic tectum of the zebrafish visual system is densely interspersed by OPCs, where they sculpt the remodelling of retinal ganglion cell (RGC) axons during phases when activity-dependent tuning occurs ^3,13–17^. To get a better sense of their proximity to RGC synapses, we used sparse labelling of individual RGCs and their pre-synapses using transgenic and knock-in fluorescent markers. In a stable transgenic line labelling all OPCs processes, we saw that they densely surround RGC terminals with intersections at both pre-synaptic or non-synaptic sites (**Fig. 1a, Supplementary Fig. 1, Supplementary Movie 1**). Live cell imaging and proximity analysis revealed that about one quarter of pre-synapses were contacted by an OPC process (**Supplementary Fig. 1b, c)**. However, when assessing where along the RGC arbour OPCs intersect, we saw that about 75% of intersections occurred at a pre-synapse (**Fig. 1b**). OPCs are highly dynamic cells that constantly extend and retract their processes within minutes (**Fig. 1c**) ^18–20^. However, timelapse imaging showed that 96% of contacts between OPCs and RGC pre-synapses remained stable over periods of at least one hour (**Fig. 1c, d**). These observations are consistent with the idea that OPCs might stabilise RGC pre-synapses.

**Figure 1:**
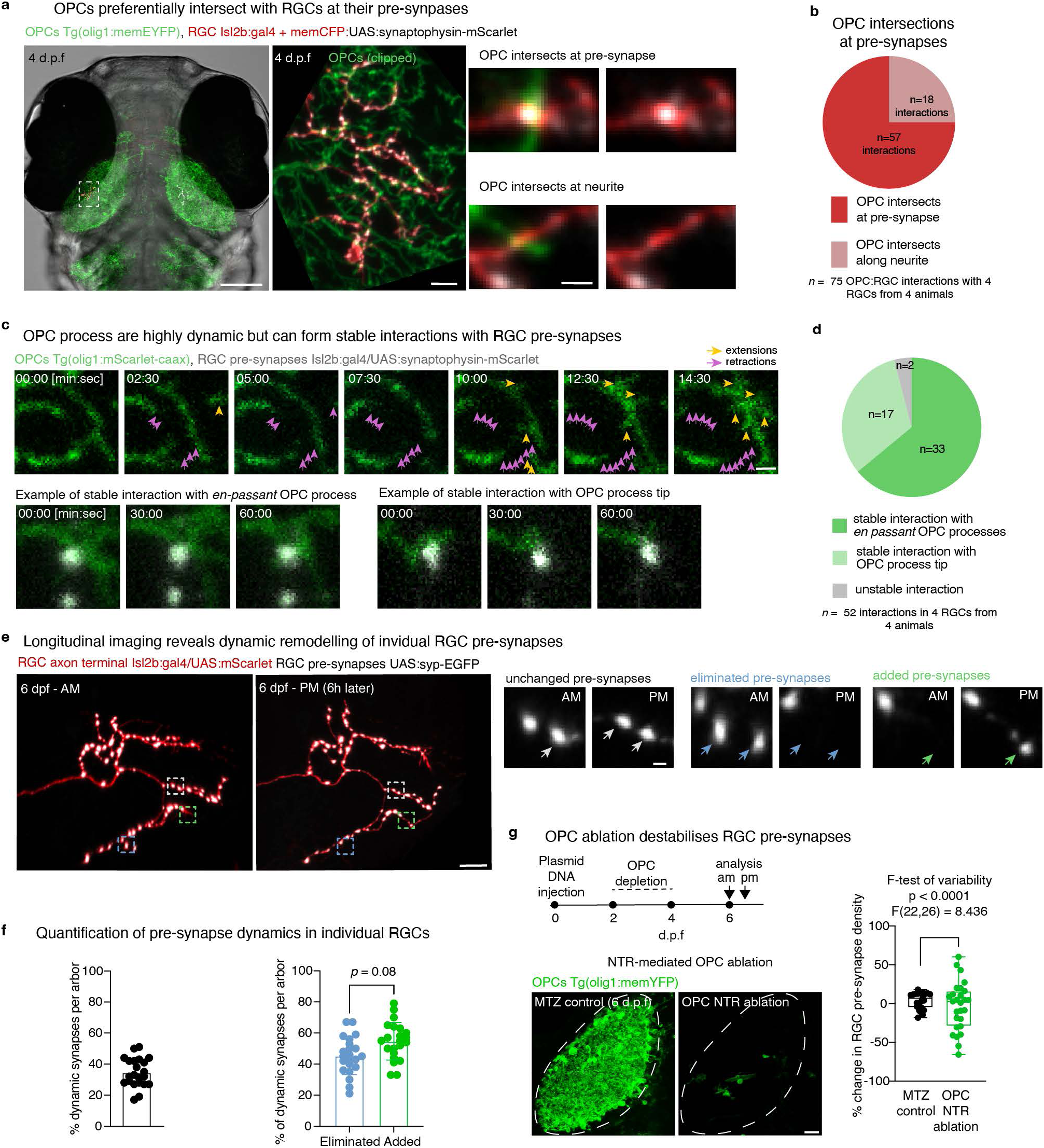
Oligodendrocyte precursor cells (OPCs) interact with and stabilise presynapses of retinal ganglion cells (RGCs) **a)** Left: overview of transgenic zebrafish labelling all OPCs processes co-labelled with a single RGC and its pre-synapses. Middle: 15µm z-projection showing a higher power view of RGC axonal arbour from animal in left (dashed box) and the surrounding OPC processes (image rotated by 90 degrees). Right: examples of OPC processes intersecting the RGC terminal at a pre-synapse and at a neurite. Scale bars: 100 µm (left), 5 µm (centre) and 1 µm (zoom-ins). **b)** Left: pie chart showing quantifications of OPC intersections with pre-synapses or neurite (n= 75 OPC:RGC interactions in 4 animals). Right: pie chart showing quantification of the number of RGC pre-synapses opposed by an OPC process (n= 232 pre-synapses in 4 animals). **c)** Time-lapse showing that OPC processes are highly dynamic but form stable contacts with RGC pre-synapses. Arrows point to retractions (magenta) and extensions (yellow) of OPC processes. Scale bar: 1 µm. **d)** Pie chart shows frequency of stable and unstable interactions of OPC processes and RGC pre-synapses (n= 52 interactions from 4 animals, 60 min timelapses). **e)** Longitudinal imaging of a single RGC reveals that individual pre-synapses can be stable, eliminated and added over a 6hr period. Dashed box indicates position of zoom-ins. Scale bars, 10 µm (left images), 1 µm (synapse zoom-ins). **f)** Left: quantification of the number of dynamic pre-synapses per RGC arbour (mean 34.55 ± 9.23 s.d). Right: quantification of the proportion of dynamic synapses that are eliminated or added per RGC arbour (mean 45.32 ± 12.05 s.d. are eliminated vs. 54.73 ± 12.08 are added, paired two-tailed t-test, t=1.83, df=21). n = 22 RGCs from 20 animals (four independent experiments). **g)** Schematic shows timeline of OPC ablation experiment. Representative images showing OPCs in metronidazole (MTZ) treated, non-NTR expressing control tectum vs. NTR-mediated ablated tectum. Dashed line indicates tectal neuropil Scale bar, 20 µm. Quantification of the change in pre-synapse density per RGC arbour within 6h plotted. (median 6.93±436.36 s.d. in control vs. 2.67±126.0 in OPC NTR ablated; Mann-Whitney U-test, U=259. F-test of variance, F(22,26)=8.44, p<0.001). n = 23/27 RGCs from 20/22 animals in control/OPC ablation, respectively (7 independent experiments).

To study the dynamics of RGC pre-synapses, we carried out longitudinal *in vivo* imaging of individual RGC terminals and their synapses at 6 days post fertilisation (d.p.f.). Over a 6-hour period, mapping the fates of 1683 individual synapses revealed that they were either stable, or dynamically changed by being eliminated or newly added (**Fig. 1e, Supplementary Fig 1g)**. These dynamic synapses accounted for approximately 35% of the total pre-synapses observed during 6-hour period (**Fig. 1f**). As synapse additions and eliminations occurred at a similar frequency, these dynamic changes only led to minor net change in overall synapse density (**Fig. 1f, g**). To test whether OPCs contribute to this stability, we eliminated them using a previously established ablation model ^3^. As hypothesised, the change in pre-synapse density for individual arbours became highly variable, with some arbours exhibiting > 50% increase or decrease in synapse density, respectively (**Fig. 1g**). These changes in synapse density following OPC ablation show that RGC pre-synapses are destabilised in the absence of OPCs.

Next, we wanted to test if RGC pre-synapse stabilisation by OPCs was through an activity-dependent feedback mechanism. It was previously shown that OPCs express machinery required for vesicular release, including Vesicle Associated Membrane Proteins 2 and 3 (Vamp2/3) ^21–23^, for which we find robust expression in our single cell RNA sequencing data of zebrafish OPCs that we expanded with additional cells (**Supplementary Fig. 2a**). We generated new transgenic reporter animals that selectively express pH-sensitive EGFP (pHluorins) fused to Vamp2/3 in OPCs, allowing study of these events by dynamic live cell imaging **(Fig. 2a, b, Supplementary Fig. 2**). Timelapse imaging revealed the presence of distinct, focal fluorescent fluctuations that were sensitive to botulinum toxin (BoNT) (**Fig. 2b, Supplementary Fig 2g, Supplementary Movie 2**). These events could either be sustained or transient (**Fig. 2b, Supplementary Fig 2b, Supplementary Movie 3**). Using an automated detection pipeline that we developed (**Supplementary Fig. 2d-f**), we saw that the frequency of OPC vesicle release varied between brain regions, being consistently higher in regions of higher spontaneous neuronal activity (**Fig. 2c, d, Supplementary Movie 4**). This suggests a positive correlation between the activity of neurons and that of OPC vesicular release.

**Figure 2:**
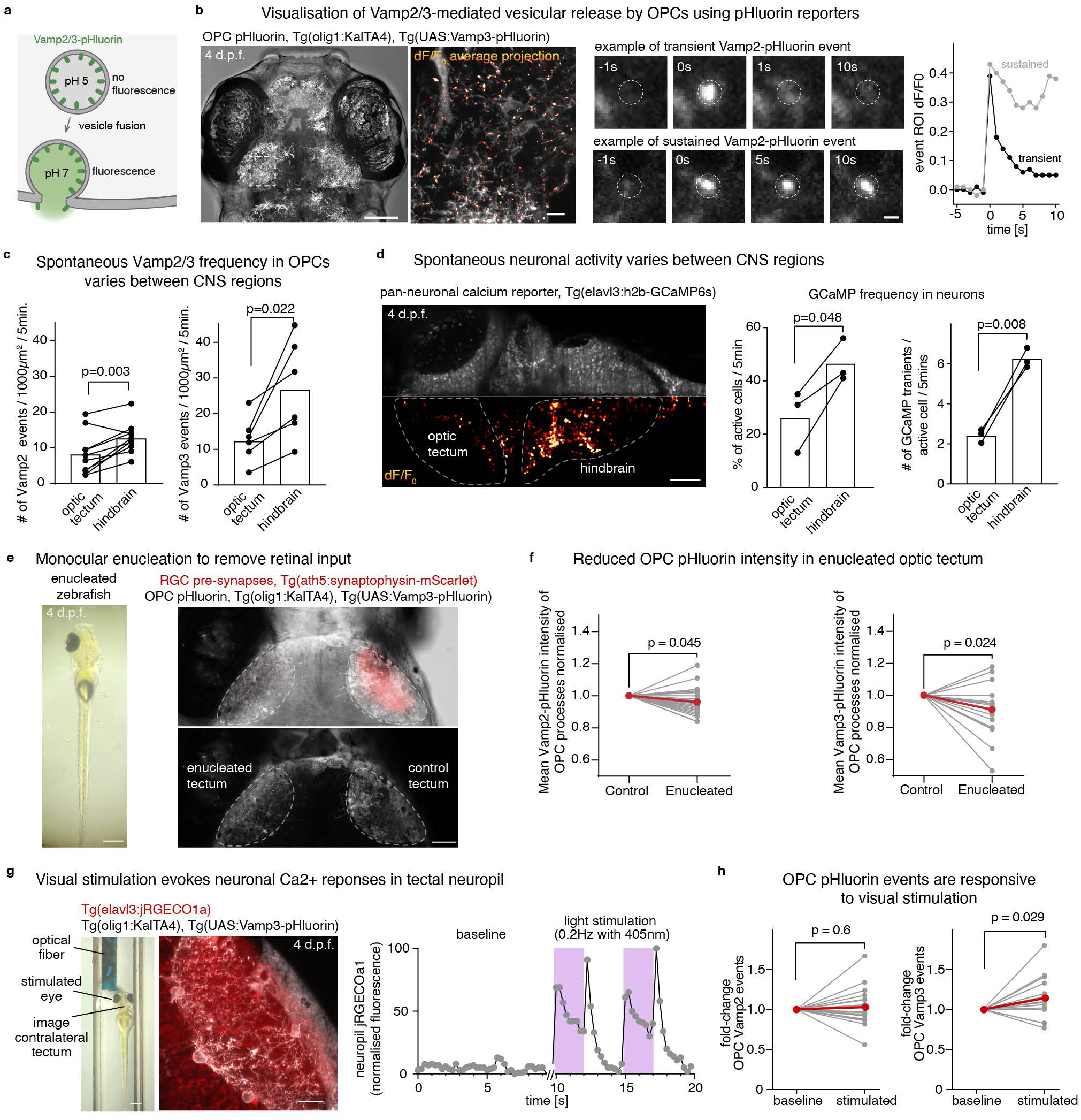
OPCs exhibit activity-dependent Vamp2/3-mediated vesicular release. **a)** Schematic of super-ecliptic pHluorin fused to Vamp2 or Vamp3 for fluorescent visualisation of vesicle release. **b)** Left: overview of transgenic zebrafish with OPCs expressing pHluorin fused to Vamp3. Middle; average projection of a 2-minutes Vamp3-pHluorin timelapse in the optic tectum overlayed with dF/F_0_ average projection illustrating fluorescence fluctuation hotspots. Right: example time-lapses showing transient (top) and sustained (bottom) Vamp2-pHluorin events. The dF/F_0_ curves of the transient (black) and sustained (grey) events were calculated using mean pixel value of a 2 µm^2^ region centred around the event point of origin. Scale bars: 100 µm (left), 10 µm (middle),1 µm (right). **c)** Quantifications showing increased spontaneous Vamp2- and Vamp3 OPC pHluorin frequencies in the hindbrain compared to the optic tectum (paired two-tailed t-tests, t=3.97, df=9 for Vamp2, t=3.27, df=5 for Vamp3). n=10/6 animals in Vamp2/Vamp3, respectively (2 independent experiments each). **d)** Left: average projection (top) and dF/F_0_ (bottom) of a 5-minute single-plane timelapse showing levels of spontaneous calcium signalling activity in neurons expressing nuclear GCaMP6s. Quantifications show increased percentage of neurons eliciting at least one Ca^2+^ transient across brain regions (left graph, paired two-tailed t-test, t=4.39, df=2) and increased frequency of Ca^2+^ transients per active cell (right graph, paired two-tailed t-test, t=10.98, df=2). n=3 animals (1 independent experiment). Scale bar: 25 µm. **e)** Left: photo of a monocularly enucleated zebrafish. Right: example confocal image illustrating that unilateral enucleation removes retinal input in the contralateral optic tectum. Scale bars: 500µm (left), 50µm (right). **f)** Quantification of the mean fluorescence intensity of Vamp2 and Vamp3-pHluorin in OPC processes in control and enucleated tectal neuropils at 4 d.p.f. (paired two-tailed t-tests, t=2.12, df=22 for Vamp2, t=2.47, df=18 for Vamp3). n=23/19 animals for Vamp2/3, respectively (2 independent experiments). **g)** Left: Image showing setup for monocular visual stimulation of zebrafish using an optic fibre with simultaneous imaging of neuronal Ca^2+^ and OPC Vamp3-phluorin activity in the contralateral optic tectum. Scale bars: 500 µm (left), 20 µm (right). Right: example traces showing ‘On’ and ‘Off’ neuronal Ca^2+^ responses in the tectal neuropil upon stimulation with 405nm light pulses at 2Hz (purple boxes). **h)** Quantification of OPC Vamp2 and Vamp3 pHluorin activity in the optic tectum at baseline and during visual stimulation at 0.2Hz (paired two-tailed t-tests, t=0.49, df=17 for Vamp2, t=2.4, df=16 for Vamp3). n=18/17 animals for Vamp2/Vamp3, respectively (3/4 independent experiments).

To test if the rate of OPC vesicular release in the optic tectum is modulated by neuronal activity, we removed retinal input through unilateral enucleation, taking advantage of the circumstance that all RGCs project to contralateral tectum in zebrafish (**Fig. 2e**) ^24^. In these animals, overall intensity of both Vamp2 and Vamp3 pHluorin were reduced in the non-innervated tectum when compared to the innervated contralateral counterpart, suggesting reduced cumulative vesicle fusion in the absence of retinal input (**Fig. 2f**). Conversely, using a visual stimulation paradigm which elicited enhanced neuronal calcium (Ca^2+^) responses in the tectal neuropil (**Fig. 2g, Supplementary Movie 5**), we found significantly enhanced Vamp3-pHluorin event frequencies in OPCs during phases of visual stimulation (**Fig. 2h**). Together, these gain- and loss-of-function manipulations indicate that OPC vesicular release rates are influenced by neuronal activity, and could therefore be a means of activity-dependent feedback to stabilise RGC pre-synapses.

To test if dynamic vesicular release by OPCs affects RGC synapse stability we generated transgenic lines in which OPCs selectively express BoNT (OPC:BoNT), reducing Vamp2/3 pHluorin intensity (**Fig. 3a, b. Supplementary Fig. 2g-h**). Longitudinal imaging of individual synapses revealed that RGC pre-synapse stability was reduced in OPC:BoNT animals (**Fig. 3c**). Closer analysis of pre-synapse dynamics showed increased eliminations and fewer additions upon blockade of OPC vesicle release (**Fig. 3c**). It is established from neuronal studies that Vamp2-mediated vesicle fusion is mediated by rises in intracellular Ca^2+^ as neurons fire ^25^. Therefore, we tested if blunting Ca^2+^ signalling in OPCs also alters the stability of RGC pre-synapses as blockade of vesicular fusion does. To do so, we generated new transgenic animals expressing the genetically encoded Ca^2+^ scavenger SpiCee^26^ specifically in OPCs, leading to a robust reduction in their Ca^2+^ signalling activity (**Fig. 3d-f, Supplementary Fig 3a**). In these animals, we did not observe any obvious changes in OPC morphology, and RGC pre-synapses were intersected by OPC processes at similar rates as in wildtypes (**Fig. 3g, Supplementary Fig. 3b**). However, overall pre-synapse density per RGC was decreased (**Fig. 3g, h**). Furthermore, longitudinal analysis revealed that RGC pre-synapse stability was reduced in animals with SpiCee-silenced OPCs, showing fewer additions and more eliminations, similar to our results where OPC vesicle release was blocked (**Fig. 3c, i**).

**Figure 3:**
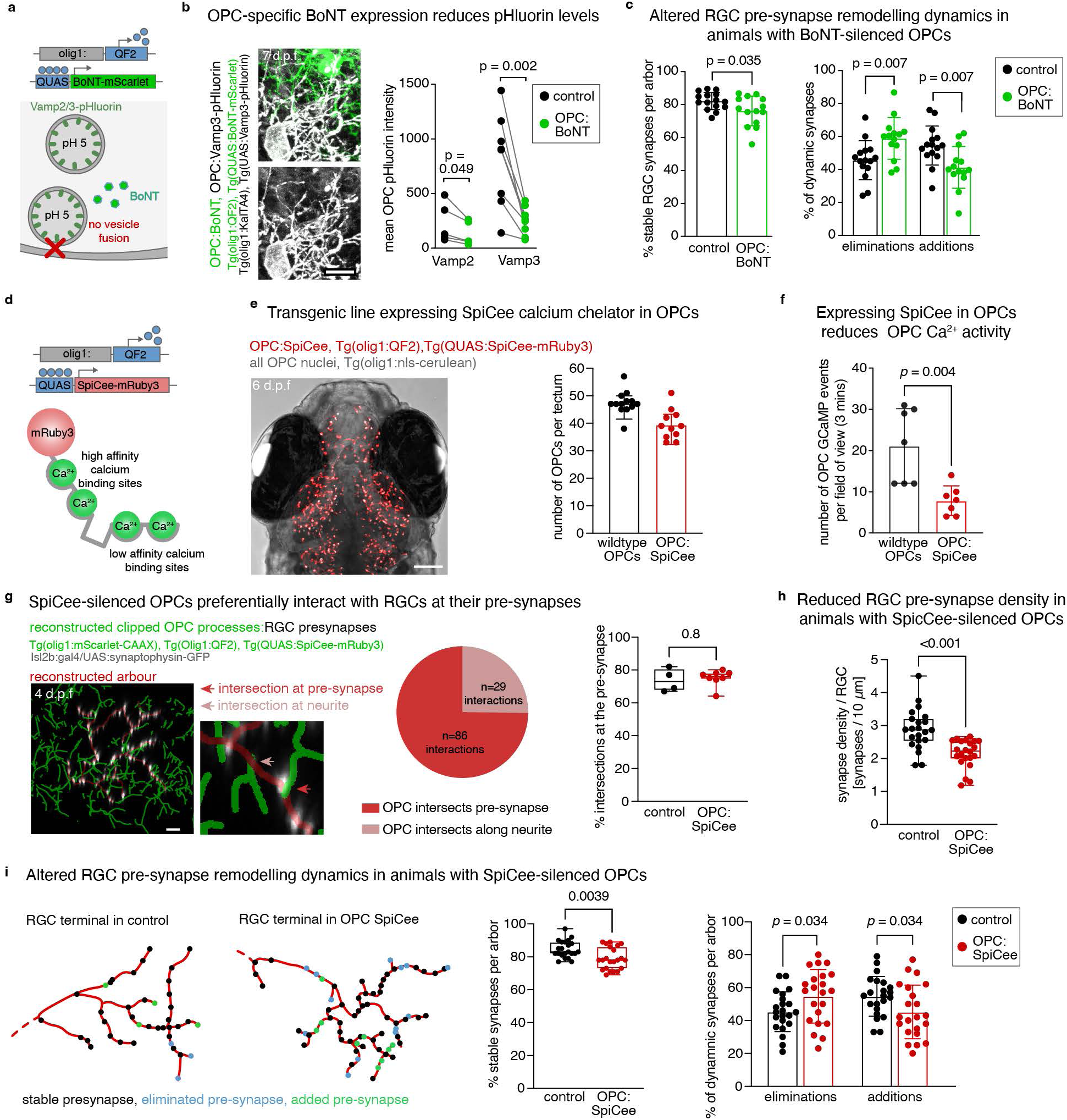
Silencing OPCs by blockade of vesicular release and dampening of Ca2+ signalling reduces RGC pre-synapse stability. **a)** Schematic of the strategy for expression of botulinum toxin (BoNT) in OPCs to block Vamp2/3-mediated vesicular release. **b)** Example image showing diminished Vamp3-pHluorin fluorescence intensity in OPCs co-expressing BoNT. Quantification shows mean Vamp2/3-pHluorin intensity of BoNT-positive and - negative OPCs within the same field of view (paired two-tailed t-tests, t=2.79, df=4 for Vamp2, t=4.71, df=7 for Vamp3). Vamp2: n=5 images from 3 animals (5-6 d.p.f. with 48 cells for control and 13 cells in BoNT group); Vamp3: n=8 images from 5 animals (5-7 d.p.f. with 133 cells for control and 25 cells in BoNT group). 1 independent experiment. Scale bar: 10 µm. **c)** Quantifications of RGC pre-synapse remodelling within a 6-hour period at 6 d.p.f. in controls and animals expressing BoNT in OPCs. Left: percentage of stable synapses (mean 82.16 ± 5.24 s.d. in control vs. 76.10 ± 9.08 s.d. OPC:BoNT, un-paired two-tailed t-test, t=2.22, df=27). Right: relative proportion of synapses that were eliminated (mean 45.55 ± 11.85 s.d. in control vs. 58.78 ± 12.64 s.d. OPC:BoNT, un-paired two-tailed t-test, t=2.91, df=27) and added (mean 54.45 ± 11.85 s.d. in control vs. 41.22 ± 12.64 s.d. OPC:BoNT, un-paired two-tailed t-test, t=2.91, df=27). n= 15/14 arbours in 9/11 animals in control/BoNT, respectively (3 independent experiments). **d)** Schematic of the strategy for expression of SpiCee in OPCs to reduce Ca^2+^ signalling activity. **e)** Left: overview of OPC:SpiCee line (using the QF2/QUAS transactivation system) co-labelled with all OPC nuclei showing broad expression of the SpiCee construct. Right: quantification shows OPC numbers at 4 d.p.f (mean 47.38 ± 4.23 s.d. in control vs. 39.45 ± 5.48 in OPC:SpiCee). n = 13/11 animals for control/OPC:SpiCee, respectively (1 independent experiment). Scale bar: 100 µm. **f)** Quantification of number of Ca^2+^ events in a Tg(olig1:GCaMP6m reporter line) in the spinal cord (mean 21.14 ± 9.03 s.d. events in control vs. 7.86 ± 3.58 in SpiCee-expressing OPCs, un-paired two-tailed t-test, t=3.62, df=12). n = 7 animals (2 independent experiments). **g)** Left: reconstructions of OPC to RGC pre-synapse interactions in OPC:SpiCee animals. Middle: pie chart shows quantifications of OPC intersections with pre-synapses or neurite. n= 115 interactions of OPC processes with 8 RGCs in 8 animals (2 independent experiments). Right: bar chart shows the number of OPC to pre-synapse intersections per arbour in wildtypes (see also Fig. 1b) and OPC:SpiCee animals (median 73.00 ± 67.50/80.75 IQR in control versus 76.00 ± 74.25/78.00 in OPC:SpiCee, two-tailed MannWhitney U-test, U = 14.50). n = 4/8 axons in 4/8 animals (2 independent experiments). **h)** Quantification showing reduced density of RGC pre-synapses in OPC:SpiCee animals (median 2.87 ± 2.52/3.21 IQR in control versus 2.23 ± 1.99/2.53 in OPC:SpiCee, two-tailed MannWhitney U-test, U = 71, p<0.001). n = 22/22 axons in 20/21 animals in control/OPC:SpiCee, respectively (4 independent experiments). **i)** Left: reconstructions of pre-synapse remodelling within a 6-hour period at 6 d.p.f. in individual RGCs in controls and OPC:SpiCee animals reveals stable, eliminated and added pre-synapses. Middle: quantification of the percentage of stable synapses (median 83.0 ± 81.75/89.00 IQR in control vs. 78.00 ± 72.75/86.25 IQR in OPC:SpiCee two-tailed MannWhitney U-test, U = 121.5, p=0.0039). Right: quantification of the relative proportion of dynamic synapses that were eliminated (mean 45.32 ± 12.05 s.d. in control vs. 54.77 ± 16.29 s.d. in OPC:SpiCee, un-paired two-tailed t-test, t=2.19, df=42) and added (mean 54.73 ± 12.08 s.d. in control vs. 45.23 ± 16.29 s.d. in OPC:SpioCee, un-paired two-tailed t-test, t=2.19, df=42). n= 22/22 arbors from 20/21 animals in control/OPC:SpiCee, respectively (4 independent experiments).

Together, our data demonstrate that the dynamic remodelling of pre-synapses is tuned by an activity-dependent vesicle release mechanism through OPCs. Several open questions remain to be addressed. Most notably, it will be interesting to address the signalling mechanisms involved by the OPC feedback pathway. What do they exocytose? Gliotransmitter release has been reported for several cell types, including activity-dependent release of GABA from OPCs ^21^. However, we do not find the expression of *gad1* required for GABA synthesis in OPCs ^20^. A recent study analysing the secretome of OPCs reports the release of many components that could directly or indirectly affect synapse stability ^27^. Notably, OPCs are major sources of extracellular matrix components and modifying enzymes, which have established roles in regulating axon and synapse development and maturation, including the opening and closure of periods of plasticity ^28,29^. Whether these are released by dynamic and BoNT-sensitive pathways remains to be investigated in future studies. However, such a scenario would place OPCs as a key player in regulating circuit plasticity in the visual system and beyond.

## Acknowledgements

We thank Dr David Lyons, Dr David McLean, and all members of the Czopka group for feedback on manuscript. We are grateful to the following individuals for sharing plasmids used in this study: Dr Rafael Almeida (University of Edinburgh) for sharing plasmids for Vamp2/3-pHluorin and SpiCee-mRuby3. Dr Marnie Halpern (Dartmouth College) for the pME_QF2 and QUAS:mApple-CAAX plasmids. Part of the computations/data handling were enabled by resources provided by the National Academic Infrastructure for Supercomputing in Sweden (NAISS), partially funded by the Swedish Research Council through grant agreement no. 2022-06725. Work in GCB lab was supported by the European Research Council (Horizon Europe Research and Innovation Programme/ ERC Advanced grant SingleMS, 101096064) and Strategic Research Area Stem Cells and Regenerative Medicine (Karolinska Institutet). Work in TC lab was funded by the European Research Council (Horizon 2020 ERC StG grant MecMy, 714440), the Wellcome Trust (Senior Research Fellowship, 224497/Z/21/Z) and the BBSRC (20RM3, BB/V017012/1 to TC). ED was supported by an EastBIO DTP studentship from the BBSRC.

## Author contributions

Conceptualization: TC; Methodology: ED, DY, PB, LJH, YX; Investigation: ED, DY, PB, LJH, YX; Formal analysis: ED, DY, EA, PB, LJH; Supervision: GCB and TC; Visualization: ED, DY, TC; Writing: ED, DY, TC; Funding acquisition: GCB and TC.

---

**Supplementary Figure 1:**
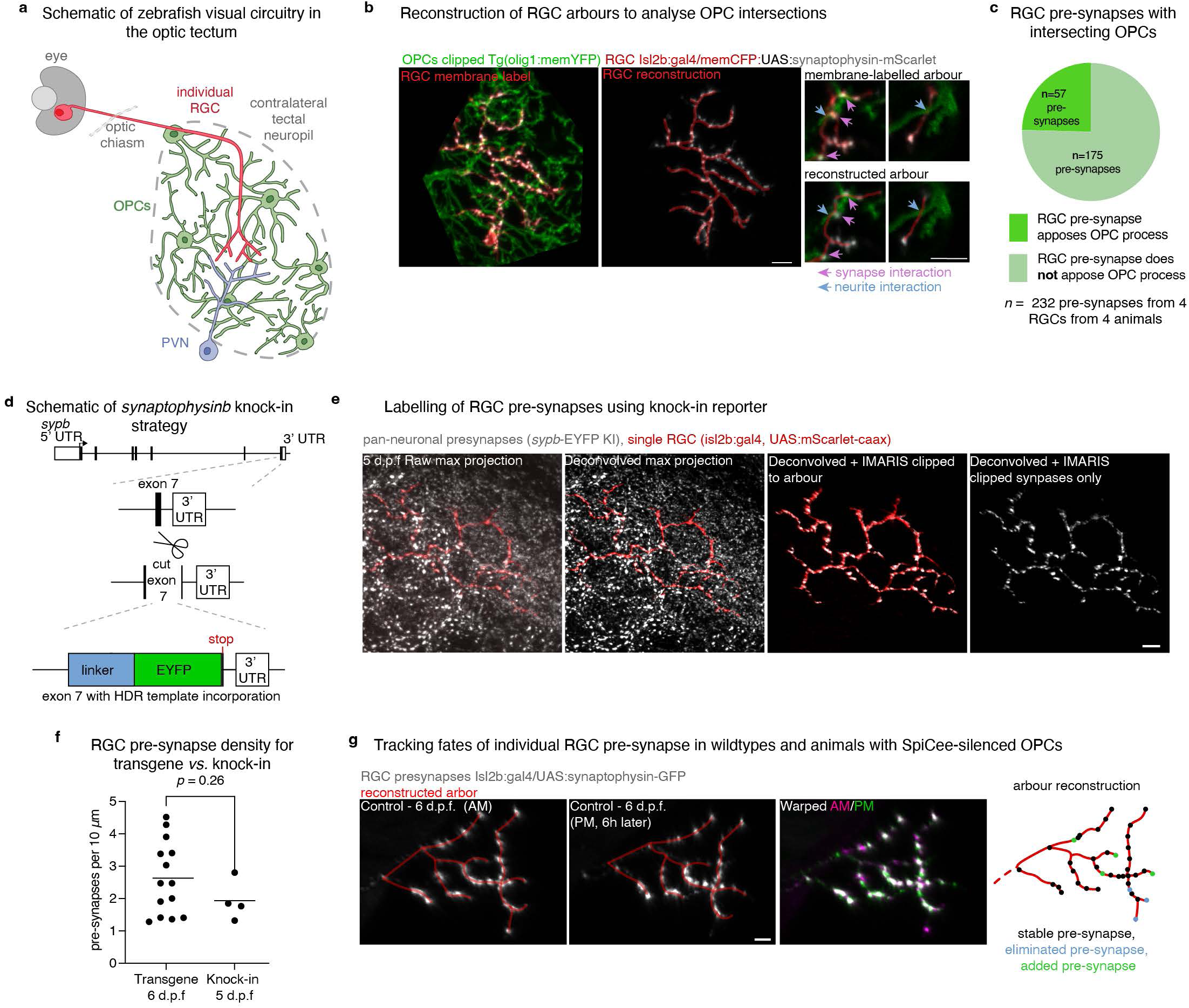
Labelling of retinal ganglion cell (RGC) pre-synapses in zebrafish and analysis of interactions with oligodendrocyte precursor cells (OPCs). **a)** Simplified schematic of the zebrafish optic tectum delineating innervation by RGC axons, relaying periventricular neurons (PVN) and the presence of OPCs. **b)** Left: confocal image of an Individual RGC terminal and its pre-synapses labelled in a full transgenic animal labelling all OPC processes. OPCs were clipped using Imaris within approx. 15 µm of the RGC terminal. Middle and right: the Imaris filament tracer (middle panel, red lines) can be used to connect RGC pre-synapses to reconstruct the arbour to analyse intersections with OPC processes. Scale bar: 5µm **c)** Pie chart showing quantification of the number of RGC pre-synapses opposed by a wild-type OPC process. n= 232 pre-synapses in 4 RGCs in 4 animals (2 independent experiments). **d)** Simplified schematic of *sypb*-EYFP KI c-terminal knock-in (KI) strategy. GuideRNA (gRNA) was designed to target the c-terminus of exon 7 of *synaptophysinb* (*sypb*) gene KI strategy, just 5’ to the stop codon. The homology-directed repair template contains a (GGGGS)_2_ linker peptide sequence and EYFP flanked with homology arms to the locus around the CRISPR cut site, to ensure in-frame c-terminus knock-in of linker-EYFP into *sypb*. **e)** Individual RGC terminal labelled in a *sypb*-EYFP-KI reporter animal. Images left to right represent workflow from raw image through deconvolution and 3D clipping to identify pre-synapses localised to the labelled RGC terminal. Scale bar: 5 µm. **f)** Quantification of pre-synapse density of single RGCs in plasmid DNA transgenic labelling (6 d.p.f) vs. knock-in synapse labelling (5 d.p.f) (mean 2.63 ± 1.13 s.d from transgenic, vs. mean 1.93 ± 0.62 s.d. from knock-in, paired two-tailed t-test, t=1.165, df=16). n= 14/4 RGC in 10/3 animals transgenic/knock-in, respectively (3/1 independent experiments). **g)** Example confocal images of pre-synapses of individual RGCs with their arbourisations reconstructed at different points in time. Overlay of timepoints (using bUnwarpJ plugin on FIJI) allows tracking the fates of individual pre-synapses as shown in reconstruction. Scale bar: 5µm.

**Supplementary Figure 2:**
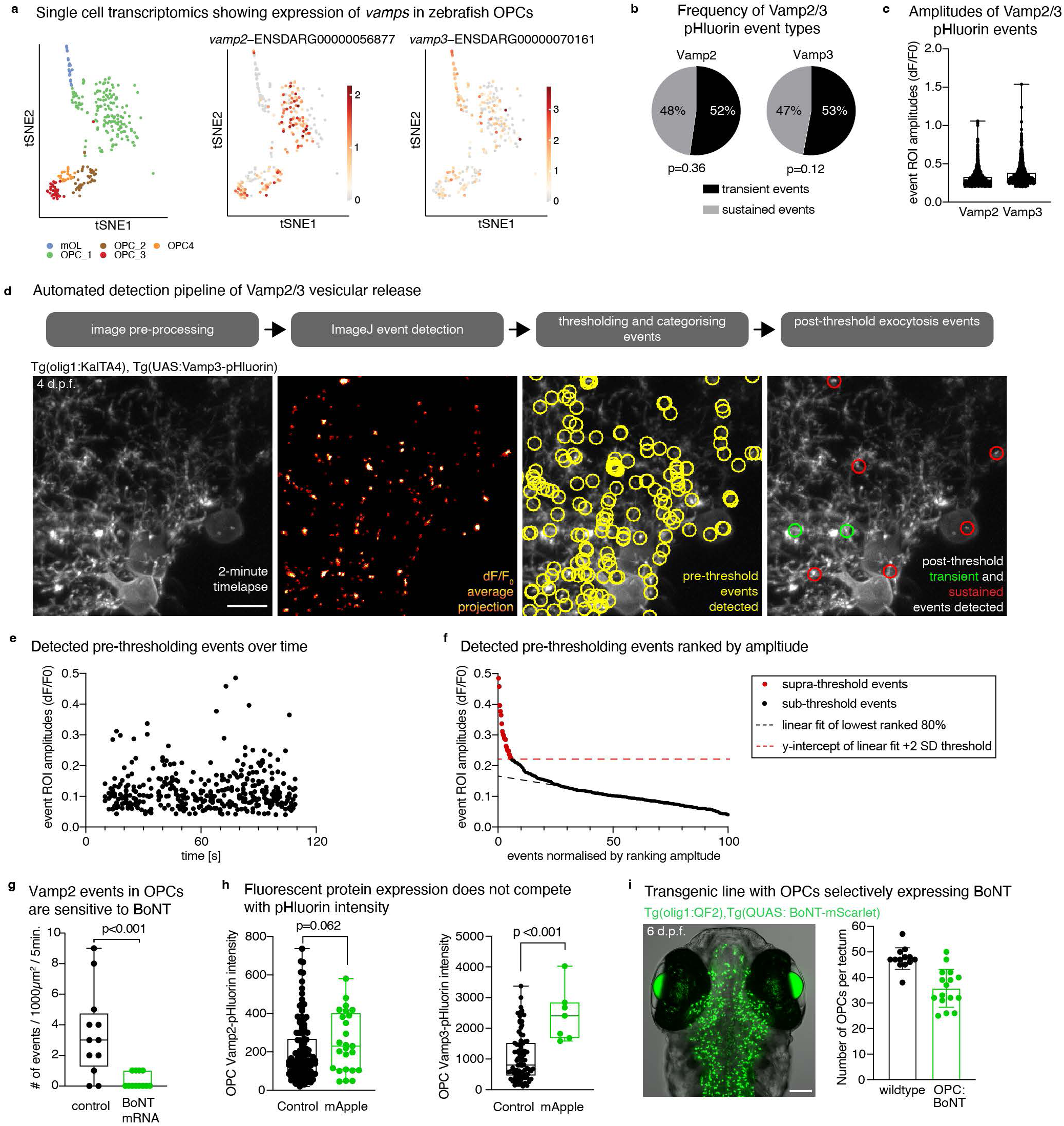
Characterisation of Vamp2/3-pHluorin signalling OPCs and the effect of BoNT. **a)** t-SNE plots of single cell RNA sequencing data from olig1:memEYFP-sorted cells with oligodendrocyte lineage identity showing expression of *vamp2* and *vamp3*. **b)** Percentage of transient and sustained Vamp2- and Vamp3-pHluorin events in the optic tectum at 4 d.p.f. (paired two-tailed t-tests, t=0.95, df=9 for Vamp2, t=1.78, df=7 for Vamp3). n=10/8 animals for Vamp2/Vamp3 (2 independent experiments). **c)** Amplitudes of individual Vamp2- and Vamp3-pHluorin events in the optic tectum at 4 d.p.f. (median 0.27 ± 0.24/0.33 IQR for Vamp2 events vs. 0.31 ± 0.27/0.39 for Vamp3 events). n=10/8 animals in Vamp2/Vamp3, respectively (2 independent experiments). **d)** Automated pHluorin detection pipeline for quantification of transient and sustained Vamp2/3-pHluorin events in OPCs. An example 1Hz volumetric light-sheet timelapse of the optic tectum was transformed to generate dF/F_0_ (moving average) to highlight transient fluorescent signals. The putative events (yellow circles) were thresholded using custom sample-based thresholding pipeline to identify supra-threshold transient (green circles) and sustained (red circles) event. Scale bar: 10 µm. **e)** Amplitudes of all putative events during example timelapse plotted against time. **f)** Amplitudes of all putative events ranked by order of decreasing amplitudes with a linear regression fitted using the lowest ranked 80% of events. The obtained y-intercept +2 s.d. value is the sample-based thresholding value for qualifying as supra-threshold events. **g)** Quantification of Vamp2-pHluorin events detected in controls and animals with single cell mRNA injection for global expression of BoNT (median 3.00 ± 1.25/4.75 IQR events for control group vs. 0.00 ± 0.00/1.00 for the BoNT mRNA group, two-tailed Mann-Whitney U-test, U = 18) n=12/12 timelapses from 6/6 animals in contro/BoNT mRNA, respectively (2 independent experiments). **h)** Quantification shows Vamp2/3-pHluorin intensity of mApple-CAAX positive and negative individual OPCs (Vamp2: median 149.29 ± 85.38/266.70 IQR for control vs. 230.10 ± 106.50/401.00 for mApple group, two-tailed Mann-Whitney U-test, U = 1212; Vamp3: median 805.00 ± 468.80/1518.00 IQR for control vs. 2410.00 ± 1686.00/2838.00 for mApple group, two-tailed Mann-Whitney U-test, U = 64). Vamp2, n = 7 images from 4 animals (with 127 cells for control and 25 cells in BoNT group); Vamp3, n = 3 images from 3 animals (with 84 cells for control and 7 cells in BoNT group), (1 independent experiments). **i)** Overview of transgenic zebrafish with OPCs expressing BoNT. Quantification shows broad co-expression of the BoNT construct in OPCs (mean 47.38 ± 4.23 s.d. OPCs in control vs. 35.81 ± 7.42 s.d. OPCs that are BoNT-expressing). n = 13/16 animals for control/OPC:BoNT, respectively (1 independent experiment). Scale bar: 100 µm.

**Supplementary Figure 3:**
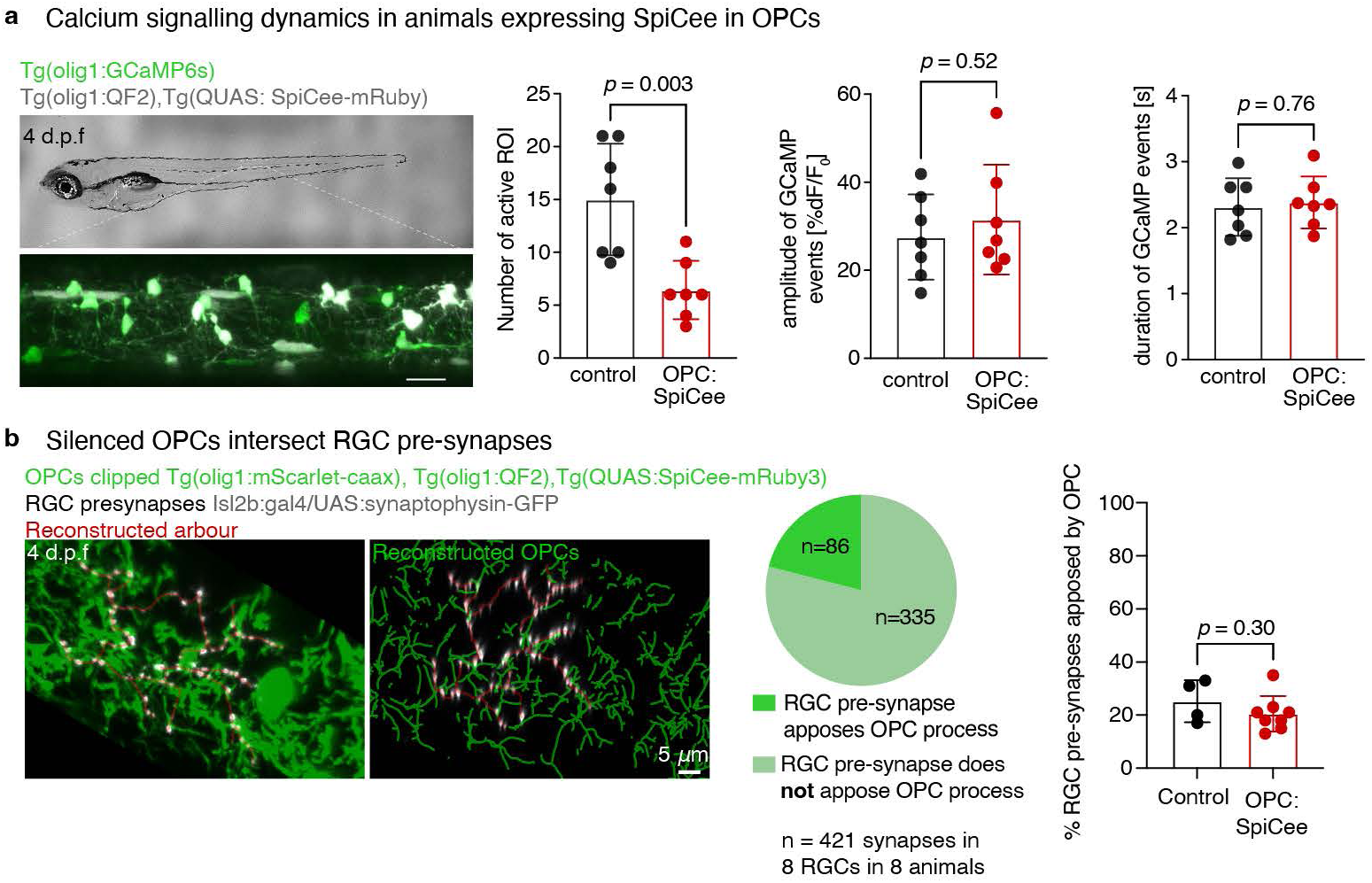
Characterisation of transgenic animals expressing SpiCee in OPCs. **a)** Left: Z-projection of confocal images of Tg(olig1:GCaMP6s), Tg(olig1:QF2,cry:CFP), Tg(QUAS:SpiCee-mRuby3) in the spinal cord. Scale bar: 20 µm. Graphs show the number active ROI detected in SpiCee expressing animals compared to non-SpiCee expressing OPC controls (left graph: mean 15.0 ± 5.3 s.d in control vs. 6.4 ± 2.8 from OPC:SpiCee, un-paired two-tailed t-test, t=3.8, df=12), the %dF/F_0_ amplitudes of detected events (middle graph: mean 27.6 ± 9.7 s.d in control vs. 31.5 ± 12.5 in OPC:SpiCee, un-paired two-tailed t-test, t=0.663, df=12), and the duration of detected events (right graph: mean 2.3 ± 0.4 s.d in control vs. 2.4 ± 0.4 in OPC:SpiCee, un-paired two-tailed t-test, t=0.311, df=12). n = 7 animals from 2 independent experiments. **b)** Left: confocal images (left) and Imaris reconstructions (right) of individual RGC arbor with its pre-synapses in a transgenic animal co-expressing SpiCee-mRuby3 (to reduce OPC Ca2+ signalling) and mScarlet-CAAX (to label the membrane of OPCs). Scale bar: 5µm. Middle: pie chart shows quantification of the number of RGC pre-synapses opposed by an OPC process. n=421 pre-synapses from 8 RGCs in 8 animals (2 independent experiments). Right: bar chart shows the proportion of RGC pre-synapses per arbour that are apposed by an OPC process in control vs. OPC:SpiCee animals (mean 25.3 ± 7.9 s.d in control vs. 20.5 ± 6.7 in OPC:SpiCee, un-paired two-tailed t-test, t=1.09, df=10). n = 4/8 RGCs from 4/ 8 animals for control/SpiCee (2 independent experiments).

**Supplementary Table 1:**
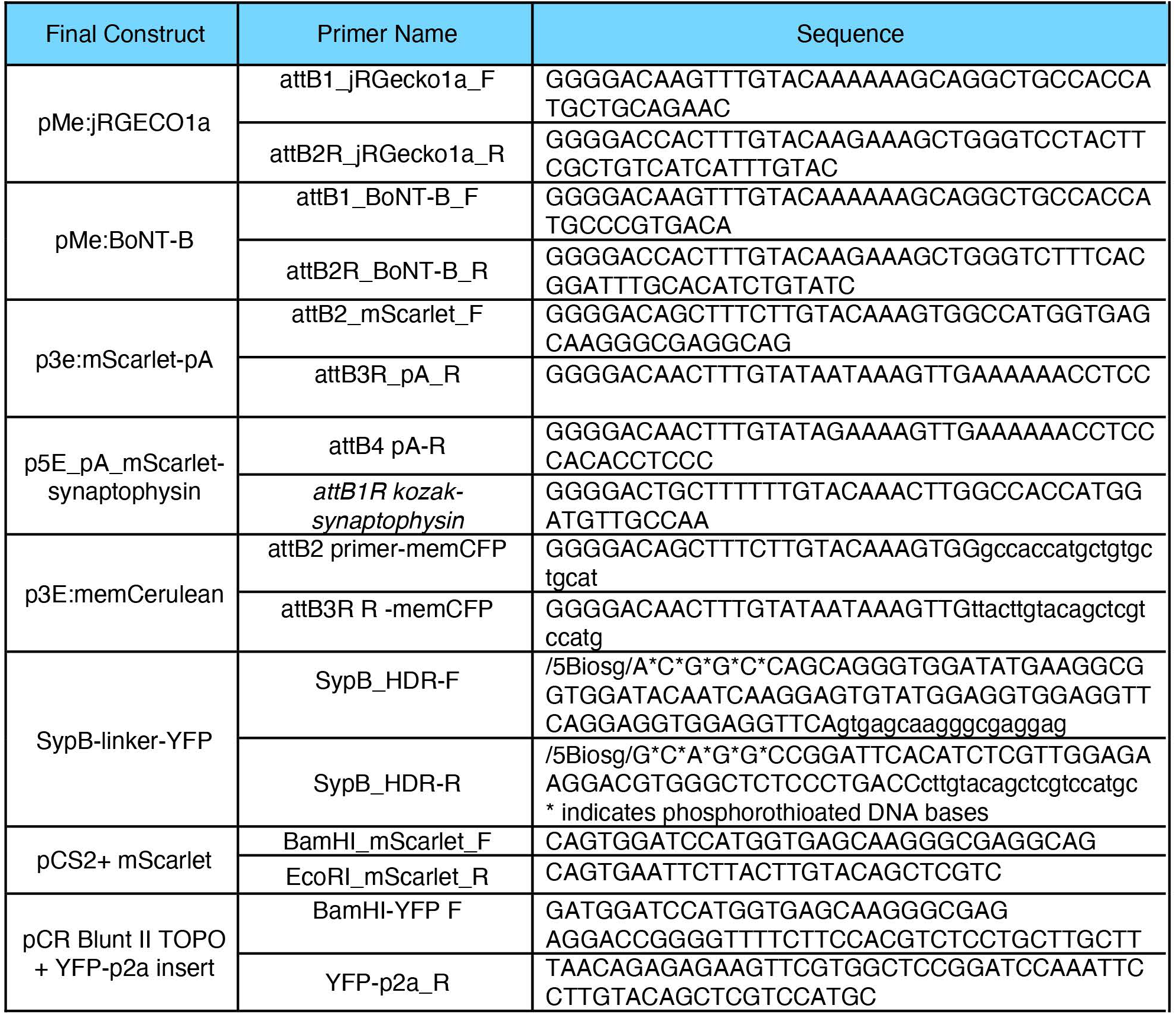
Sequences of primers used to generate zebrafish transgenesis and knock-in constructs.

**Supplementary Movie 1: Pan and Zoom of OPCs and RGCs in the optic tectum of zebrafish**

**Supplementary Movie 2: OPC Vamp3-pHluorin timelapse and dF/F_0_ transformations shown in Fig 3b. Scale bar: 10µm.**

**Supplementary Movie 3: Timelapse of a transient and sustained OPC Vamp2-pHluorin event shown in Fig 3c. Scale bar: 1µm.**

**Supplementary Movie 4: Whole brain neuronal GCaMP timelapse and dF/F_0_ transformations shown in Fig 3e. Scale bar: 20µm.**

**Supplementary Movie 5: Neuronal jRGECO1a timelapse of the optic tectum during monocular visual stimulation shown in Fig 3f,g. Scale bar: 20µm.**

## Methods

### Data and materials availability

All data underlying this study will be made available upon reasonable request. Raw sequence data, gene expression and cell type annotation tables of newly generated scRNA sequencing data of zebrafish OPCs have been deposited in the Gene Expression Omnibus (GEO) (accession number GSE313580). A browsable web resource is available at https://castelobranco.shinyapps.io/dreropc_rna_2020/

### Code availability

The code used for analysis of Ca^2+^ imaging, pHluorin intensity and pHluorin event detection is available at: 10.5281/zenodo.18196206

### Zebrafish lines and husbandry

We used the following existing zebrafish lines and strains: Tg(olig1:memEYFP)^tum107Tg 20^, Tg(olig1:mScarlet-caax)^tum111Tg 30^, Tg(olig1:CFP-NTR)^tum113Tg 3^, Tg(olig1:GCaMP6m)^tum105Tg 20^, Tg(olig1:KalTA4)^tum106Tg 20^, Tg(olig1:nls-Cerulean)^tum108tg 20^ and Tg(elavl3:h2b-GCaMP6s)^jf5Tg 31^. The following lines were newly generated for this study: Sypb-EYFP KI (knock-in), Tg(ath5:synaptophysin-mScarlet), Tg(elavl3:jRGECO1a), Tg(olig1:mScarlet), Tg(olig1:QF2), Tg(QUAS:SpiCee-mRuby3), Tg(QUAS:BoNT-mScarlet), Tg(UAS:Vamp2-pHluorin), and Tg(UAS:Vamp3-pHluorin). All animals were kept at 28.5°C with a 14-h/10-h light/dark cycle according to local welfare regulations for animals at protected stages. All experiments carried out with zebrafish at protected stages were approved by the Animals in Science Regulation Unit of the UK Home Office (PP4944080 to TC), or by the government of Upper Bavaria (Regierung Oberbayern - Sachgebiet 54; ROB-55.2-1-54-2532.Vet_02-18-153, ROB-55.2-2532.Vet_02-15-199, and ROB-55.2-2532.Vet_02-15-200 to TC).

### DNA constructs for zebrafish transgenesis and knock-ins

Sequences for all primers used are listed in **Supplementary Table 1**. We generated the following entry clones for Gateway recombination compatible with the Tol2Kit ^32^ by PCR amplification from template plasmids containing the respective coding sequences: pME:jRGECO1a, pME:BoNT-B, p3e:mScarlet-pA, p5E_pA_mScarlet-synaptophysin and p3E:memCerulean. Sequence specific primers contained appropriate *att* recombination sites for recombination with pDONR221, pDONR p2R-p3 or pDONR p4 p1R using BP clonase (Thermo Fisher Scientific). The 5’ entry clone p5E_olig1(4.2) and the middle entry clones pME_mScarlet and pME_synaptophysin_nostop were previously published by us ^20,33^. The following plasmids were kind gifts from Dr Rafael Almeida (University of Edinburgh): pME_SpiCee-mRuby3 ^26,34^, p3E_linker01xpHluorin-pA ^22^, pME-(zf)vamp2-nostop ^22^, pME-(zf)vamp3-nostop ^22^, pME_10x-Janus ^35^. The p5E_ath5 ^32^ was a kind gift from Dr Caren Norden. The pME_QF2 (Addgene plasmid # 61375), p5E:QUAS (Addgene plasmid # 61374), and pTol2_QUAS:mApple-CAAX ^36^ are gifts from Dr Marnie Halpern. These plasmids were recombined with additional clones from the Tol2Kit using LR Clonase II Plus (Thermo Fisher Scientific) to generate the following Tol2 transgenesis constructs: pTol2_olig1(4.2):QF2, pTol2_QUAS:SpiCee-mRuby3, pTol2_UAS:Vamp2-pHluorin, pTol2_UAS:Vamp3-pHluorin, pTol2_elavl3:jRGECO1a, pTol2_QUAS-BoNT-mScarlet, pTol2_UAS_synaptophysin-mScarlet, pTol2_synaptophysin-mScarlet_UAS(Janus)_memCFP. We also used the following existing constructs: pTol2_isl2b: Gal4-VP16 ^16^, pTol2_UAS:mScarlet ^20^, pTol2_UAS:Synaptophysin-EGFP ^37^, pTol2_olig1:mScarlet ^3^.

A construct for C-terminal knock-in of EYFP in the *synaptophysin b (sypb)* locus was created following the method described in ^38^, using the following gRNA sequence to target exon 7 of *sypb*: CAATCAAGGAGTGTATGGTCAGG. A homology directed repair (HDR) template containing a (GGGGS)_2_ linker peptide and EYFP flanked by sequences complimentary to the *sypb* gRNA cut site were generated using primers listed in **Supplementary Table 1**. The HDR template was amplified off a plasmid containing EYFP containing plasmid through PCR.

### Generation of messenger RNA constructs

Messenger RNAs for BoNT, mScarlet, and Tol2 transposes were by in vitro transcription using the mMESSAGE mMACHINE™ SP6 Transcription Kit (Thermo Fisher Scientific) from the following plasmids: pCS2+_BoNT LC-B ^39^ (gift from Rafael Almeida), pCS2+_mScarlet, and pCS2+_Tol2_transposase ^32^.

### DNA microinjections for sparse labelling and generation of new lines

DNA microinjections were carried out and previously described ^3^. For the generating of knock-in lines injection mixes contained the following: 70-120 ng/µl of HDR template, DNA-PK inhibitor NU7441(Selleck), 50 ng/µl of gRNA and 5 µM of Cas9 (Thermo Fischer Scientific). ‘Tg(promoter:reporter)’ denotes a stable transgenic line, whereas ‘promoter:reporter’ alone indicates that a respective plasmid DNA was injected for sparse labelling of individual cells. ‘Gene-reporter KI’ denotes a stable knock-in line.

### BoNT mRNA microinjections for global silencing of VAMP2/3-pHluorin events

For global BoNT expression in VAMP2–pHluorin timelapse experiments, embryos were co-injected at the one-cell stage with BoNT mRNA (200 ng/μl) and mScarlet mRNA (200 ng/μl), or with mScarlet mRNA alone as a control. Animals were manually dechorionated at 2 d.p.f. and screened for high mScarlet fluorescence as a proxy for relative BoNT expression levels before experimental imaging at 4 d.p.f.

### Nitroreductase-mediated ablation of OPCs

For nitroreductase (NTR)-mediated OPC ablation at early developmental stages, Tg(olig1:CFP-NTR),Tg(olig1:memEYFP) zebrafish at 2 d.p.f. were incubated in 10 mM metronidazole (MTZ) (Sigma-Aldrich) dissolved with 0.2% DMSO in 0.3× Danieau’s solution for 48 h total at 28 °C. MTZ was refreshed on in the morning and evening of 3 d.p.f.. From 4 d.p.f. embryos were kept in 0.3× Danieau’s solution until analysis. Non-NTR-expressing zebrafish Tg(olig1:memEYFP) were treated with 10 mM MTZ were used as controls in all experiments.

### Mounting of embryonic and larval zebrafish for live cell microscopy

For confocal microscopy, zebrafish were anaesthetized and embedded in agarose as previously described ^3^. For lightsheet microscopy, zebrafish were immobilised by immersion in 1 mg/ml α-Bungarotoxin (Bio-Techne) for 2 minutes, following by re-immersion in Danieau’s. Animals were embedded in 1% low melting point agarose in a U-shaped glass capillary attached to a glass-bottom dish and immersed in Danieau’s solution.

### Confocal microscopy

Images of embedded zebrafish were taken with a Leica TCS SP8 confocal laser scanning microscope. We used 448 nm for excitation of Cerulean/tagCFP; 488 nm for EGFP/GCaMP/pHluorin; 514 nm for EYFP and 552 nm for red fluorescent proteins. For overview images we used a ×10 / 0.4 NA objective (acquisition with 283-668-nm pixel size (x–y) and 2-4μm z-spacing). For all other analyses, we acquired 8-bit or 12-bit confocal images using a ×25 / 0.95 NA water objective with 56-227nm pixel size (x–y) and 0.3-3.0-μm z-spacing. For timelapse imaging of OPC processes and RGC pre-synapse dynamics, 12-bit depth 8kHz resonance scanning confocal z-stacks were acquired using a ×25 / 0.95 NA water objective with 113 nm pixel size (x–y) and 1-μm z-spacing every 30 seconds for 1 hour.

### Lightsheet microscopy

All functional imaging was performed using 12-bit image acquisition on a Leica TCS SP8 Digital LightSheet with 488 nm excitation for GCaMP/ pHluorin, and 552 nm excitation for jRGECO1a using appropriate detection filters and a Hamamatsu Orca Flash 4.0 V3 camera. For whole-brain imaging of Tg(elavl3:h2b-GCaMP6s), a 2.5× /0.07 NA illumination objective and a 10× /0.3 NA detection objective with 5 mm deflection mirrors were used (acquisition with 360nm pixel size (x–y) at single plane). Images were acquired at a frame rate of 2 Hz for 5 minutes. All other transgenic lines imaged using lightsheet microscopy used a 2.5× /0.07 NA illumination objective and a 25× /0.3 NA detection objective with 5 mm deflection mirrors (acquisition with 143-286nm pixel size (x–y) and 2-3.5μm z-spacing unless single-plane). For GCaMP imaging in OPCs, 3-minute time-lapses of 42μm Z-stacks at the level of the spinal cord were acquired at 1Hz. For pHluorin imaging in OPCs, 34μm Z-stacks were imaged with a frame rate of 1Hz for 2-5 minutes. Z-stacks at the level of the optic tectum and the caudal hindbrain were positioned 20-54μm ventrally from the surface of the neuropil. For Tg(elavl3:jRGECO1a) single-plane imaging and visual stimulation validation, 5-minute timelapses were acquired 30μm ventral from the visible tectal neuropil at a frame rate of 4 Hz.

### Enucleation and quantification of pHluorin intensity of OPC processes

Monocular enucleation was carried out in Tg(ath5:synaptophysin-mScarlet), Tg(olig1:KalTA4), Tg(UAS:Vamp2/3-pHluorin) using a tungsten needle at 2 d.p.f. Animals were screened at 4 d.p.f. for absence of synaptophysin labelled RGC axons in the contralateral tectum to the enucleated eye. Using the confocal microscope, a 6μm Z-stack of the left and right optic tectum was acquired 30μm ventral to the visible tectal neuropil. Average projections of the Z-stack were processed using a custom FIJI script to quantify the mean pHluorin intensity of OPC processes between the control and enucleated tectum.

### Analysis of OPC-RGC pre-synapse interactions

To analyse OPC-RGC pre-synapse *vs.* RGC neurite interactions, RGC arbour terminals were reconstructed through connecting labelled pre-synapses using the filament tracer in Imaris software. RGC pre-synapses were defined as discrete-punctate accumulation of fluorescence larger than 0.5 μm in diameter. On Imaris, every z-step containing the RGC arbour was analysed and observed for OPC intersections with neurite (OPC processes touches non-synapse containing reconstruction of the arbour) or intersections with a pre-synapse (pre-synapse is touched by an OPC process). For presentation, OPC processes were reconstructed using Imaris filament tracer. The number of RGC pre-synapses directly apposed by an OPC process were classified as being co-localised with any amount of OPC process.

For analysing dynamic interactions between RGC pre-synapses and OPC processes, timelapses were taken over 1h within 30-second intervals and with 1 μm spacing. Three-dimensional (3D) movies were generated, and registered using StackReg plugin using FIJI software. Interactions were classified as one of the following: *en* passant (where an RGC pre-synapse is contacted along the length of an OPC process), or at the tip. Interactions were classified as stable when contacts were maintained for the majority of the time (i.e. not counting transient moving last up to two frames) and including the start and end of the timelapse.

### Analysis of RGC pre-synapse remodelling

Each arbour was traced using Imaris Filament tracer from its first branch point to all the branch tips through connecting the dots of the labelled pre-synapses and the measurement for filament length was extracted. For fate-mapping of individual pre-synapses, the PM image was warped onto the AM image using the bUnWarp plugin on FIJI ^40^, and the brightness of fluorescence normalised to each other (**Supplementary Fig 1f**). A pre-synapse was classified as discrete punctate accumulation of the synaptophysin label and counted for the AM and PM image. Stable pre-synapses were classified as being present in the AM and PM images, eliminated pre-synapses are present in the AM only, and added pre-synapses present in PM only.

### Analysis of the number of OPCs per optic tectum

The number of OPC soma within the left tectal neuropil 4 d.p.f were counted for controls: Tg(olig1:mScarlet) or SpiCee expressing OPCs: Tg(olig1:QF2), Tg(QUAS:SpiCee-mRuby3) or BoNT-B expressing OPCs:Tg(olig1:QF2), Tg(QUAS:BoNT-B-mScarlet).

### Quantification of pHluorin intensity of OPC soma

To assess the effects of BoNT on cumulative Vamp2/3-pHluorin activity in OPCs, each transgenic line was crossed to Tg(olig1:QF2) and injected with either pTol2_QUAS:BoNT-mScarlet (**Fig. 3b**) or pTol2_QUAS:mApple-CAAX (control for UAS/QUAS competition, **Supplementary Fig. 2h**) at single-cell stage to achieve sparse expression. Large volume confocal stacks were acquired in brain regions where BoNT- or mApple-positive OPCs were sparsely labelled, and a 12.5μm^2^ circular ROIs were manually drawn on the somas of all OPCs to extract mean fluorescent values from both pHluorin and BoNT-mScarlet/mApple channels. To distinguish BoNT/mApple-positive cells, we used a common outlier detection method based on the interquartile ranges. In brief, only cells whose mean fluorescent value for BoNT-mScarlet or mApple channel that is greater that Q3 + 1.5 × (Q3 – Q1) value were classified as belonging to the BoNT/mApple group. Zebrafish were imaged between 5-7 d.p.f..

### Visual stimulations

The reliability of the visual stimulation protocol was validated using Tg(elavl3:jRGECO1a) larvae at 4 d.p.f. Larvae were imaged on the lightsheet microscope using a custom glass-bottom dish with an optical fiber (Thorlabs) embedded within the U-shaped capillary to project light towards one eye (Fig. 3f). Using a DC2200 LED driver (Thorlabs) coupled to a 405nm LED light source (M405FP1-Thorlabs) at 0.2% output power, reliable neuronal ‘on’ and ‘off’ responses were achieved with a 0.2Hz stimulation protocol (200ms ON, 300ms OFF).

### Analysis of calcium signalling events

For the quantification of basal neuronal Ca^2+^ event frequency in the hindbrain and optic tectum, timelapse recordings were first registered in FIJI using the StackReg plugin. Using Suite2p (a Python-based package) for cell segmentation, the mean fluorescent values of all ROIs (active and inactive cells) across time within defined brain regions were extracted and analysed using a custom MATLAB code to determine Ca^2+^ transients frequency for each cell.

To assess the effect of SpiCee on OPC Ca^2+^ event frequency in Tg(olig1:GCaMP6m), maximum projections of OPC Ca^2+^ timelapses were first registered in FIJI using the StackReg plugin. ROIs were manually drawn (blinded) around areas of OPC processes that showed increased brightness associated with increased GCaMP6m signal. The mean fluorescent values across time from drawn ROIs were exported to MATLAB and converted to dF/F_0_ with a moving minimum used as baseline, to correct for bleaching effects. Event detection was done through the same MATLAB code used for neuronal event detection. The detection threshold was set as the highest value between 4SD or 10% increase from the mean fluorescence.

### Analysis of OPC Vamp2/3-pHluorin event frequency

For the analysis and quantification of Vamp2/3-pHluorin events, maximum projections of each timelapse were first registered in FIJI using the StackReg plugin. Next, the auto-fluorescence from the skin and other irrelevant signals were cropped from each frame, followed by a dF/F_0_ transformation using a 5-frame moving average as the baseline is generated to highlight transient fluorescent signals (**Supplementary Fig. 2d; Supplementary Movie 2**). The onset and the co-ordinates of each putative event was calculated using the TrackMate plugin and a custom FIJI script. A circular 2μm^2^ ROI was then marked for each putative event and the mean fluorescent values of each ROI across time is measured from the raw timelapse. dF/F_0_ curves were generated for each event using 5 frames preceding the event as baseline. Events with an irregular dF/F_0_ curve, a dF/F_0_ amplitude below 0.04, and events that began within the first/last 10 seconds of the recording were excluded. The dF/F0 amplitudes of all putative events from the recording were then ranked by order of decreasing amplitudes with a linear regression fitted using the lowest ranked 80% of events (**Supplementary Fig. 2e, f**). The obtained y-intercept +2 s.d. value is the sample-based thresholding value for qualifying as supra-threshold events. Events with transient amplitudes above 50% max dF/F_0_ after 5 seconds were categorised as sustained events. Error matrix calculations were performed using timelapses from 8 animals and reports F1 score of 0.99 (0.97 precision, 1.00 recall, 0.93 specificity, 0.98 accuracy).

To assess the effect of visual stimulation on Vamp2/3-pHluorin events, visual-stimulation was initiated after 2-minute 20-seconds in a 4-minute 20-seconds continuous timelapse recording. Total pHluorin events were assessed during the 2-minute stimulation period and the 2-minute period preceding stimulation (baseline). All putative events detected from both baseline and stimulation periods within the same timelapse underwent sample-based thresholding together.

### Single cell RNA sequencing

OPCs were isolated by fluorescent activated cell sorting exactly as described previously ^20^. Sequencing of the single cell was performed on Illumina HiSeq2500 with the following specifications: The run was on a high-output flow cell with 50 bp single-read, clustered on a cBot with V4 kit. Reads were trimmed with Cutadapt 1.8.0 and aligned to the transcriptome using STAR 2.5.1.b ^41^ to the reference genome GRCz11 and using ENSEMBL94 transcripts annotations with the following parameters: -sjdbFileChrStartEnd SJ.out.tab, -sjdbScore 0, -outFilterMatchNmin 10, - outSAMunmapped Within, -quantMode TranscriptomeSAM. Including the splicing junction database calculated by STAR on the same single cells in a first alignment. Aligned single cell reads were sorted and transformed to bam files using Samtools 1.3. Gene expression was calculated with Salmon 0.9.1 using the sorted bam files as input. From the outputs we used the TPM gene expression values to build the expression matrix as detailed in ^20^.

### Analysis and presentation of sequencing data

Cells were clustered with Seurat v3.2.1. Cells were filtered based on the distribution of gene expression (minimum 500 expressed genes per cell) and mitochondrial gene expression (maximum 0.05%). The remaining 310 cells were log-normalized individually with a scale of factor of 10,000.

For downstream analyses we used the top 2000 variable genes. The shared-nearest neighbour (SNN) graph was constructed on cell-to-cell distance matrix from top 50 PCs. The SNN graph with resolution 1 was used as an input for the smart local moving (SLM) algorithm to obtain cell clusters, and visualized with t-distributed stochastic neighbour embedding (t-SNE). The cells from the 2 sequenced plates, one from our previous study and a second plate, were merged into a single dataset using Seurat v3 merge function, leading to 510 cells. In order to integrate both datasets we used the cell annotations from ^20^ as reference to find the anchors with FindTransferAnchors function and 20 dimensions. Then we performed a label transfer to classify all the cells following our annotations from ^20^. The final dataset consists in 510 cells, 171 OPC1, 41 OPC2, 35 OPC3, 20 OPC4, 26 mOL 60 VLMC1 and 157 VLMC2. The dataset was subset to extract the OPC1 and mOL to show the expression of the candidate gene list. Plots were build using ggplot2 in R 4.0.0.

### Image and data presentation

Images were analysed with Fiji and Imaris. Morphology reconstructions were carried out with the Imaris FilamentTracer module. Images were deconvolved using Huygens. Data were prepared and assembled using GraphPad Prism, Fiji and Adobe Illustrator.

### Statistics and reproducibility

For analyses that involved cohorts of animals or treatment groups, zebrafish embryos of all conditions were derived from multiple clutches and selected at random before treatment. No additional randomization was used during data collection. For time-course and time-lapse analyses, zebrafish were screened before imaging, with appropriate labelling being the only criterium for inclusion in the experiment. Following time-lapse imaging experiments, some animals were discarded when there was too much z-drift, preventing accurate analysis. When analysing RGCs, arbours that could not be traced back to the optic nerve were excluded for analysis, as well as arbours that showed signs of ill health such as blebbing. When carried out, all exclusions were blind to the experimental condition. No other data were excluded from the analyses. Sample sizes were chosen based on similar sample sizes that were previously reported ^42–45^. No statistical analysis was used to pre-determine sample sizes. Data collection and analysis were not performed blind to the conditions of the experiments unless stated. Statistical analysis was performed using Microsoft Excel and GraphPad Prism. All data were tested for normal distribution using the Shapiro–Wilk normality test before statistical testing. Normally distributed data are shown in the graphs as mean ± standard deviation (s.d.). Non normally distributed data are shown as box and whiskers plots of the median ± interquartile ranges (I.Q.R) and minimum/ maximum values. Paired data are clearly labelled as paired data. For statistical tests of normally distributed data that compared two groups, we used paired and unpaired t-tests, as appropriate. Non-normally distributed data were tested using the Mann–Whitney U test (unpaired data).

## References

1. Holtmaat, A. & Svoboda, K. Nat. Rev. Neurosci. 10, 647–658 (2009).

2. Barabási, D.L., Castro, A.F. & Engert, F. Nat. Rev. Neurosci. 26, 232–243 (2025).

3. Xiao, Y., Petrucco, L., Hoodless, L.J., Portugues, R. & Czopka, T. Nat. Neurosci. 25, 280–284 (2022).

4. Auguste, Y.S.S. et al. Nat Neurosci 25, 1273–1278 (2022).

5. Buchanan, J. et al. Proc National Acad Sci 119, e2202580119 (2022).

6. Xiao, Y. & Czopka, T. Nat. Neurosci. 26, 1663–1669 (2023).

7. Chung, W.-S., Baldwin, K.T. & Allen, N.J. Cold Spring Harb. Perspect. Biol. 16, a041352 (2024).

8. Schafer, D.P., Stevens, B., Bennett, M.L. & Bennett, F.C. Cold Spring Harb. Perspect. Biol. a041810 (2024).doi:10.1101/cshperspect.a041810

9. Hill, R.A., Nishiyama, A. & Hughes, E.G. Cold Spring Harb. Perspect. Biol. a041425 (2023).doi:10.1101/cshperspect.a041425

10. Gallo, V., Mangin, J.-M., Kukley, M. & Dietrich, D. The Journal of Physiology 586, 3767–3781 (2008).

11. Bergles, D.E., Jabs, R. & Steinhäuser, C. Brain Res Rev 63, 130–137 (2010).

12. Maldonado, P.P. & Angulo, M.C. Neurosci 21, 266–276 (2015).

13. Hua, J.Y. & Smith, S.J. Nature Neuroscience 7, 327–332 (2004).

14. Hua, J.Y., Smear, M.C., Baier, H. & Smith, S.J. Nature 434, 1022–1026 (2005).

15. Smear, M.C. et al. Neuron 53, 65–77 (2007).

16. Fredj, N.B. et al. J Neurosci 30, 10939–10951 (2010).

17. Kutsarova, E., Munz, M. & Ruthazer, E.S. Front. Neural Circuits 10, 111 (2017).

18. Kirby, B.B. et al. Nature Neuroscience 9, 1506–1511 (2006).

19. Hughes, E.G., Kang, S.H., Fukaya, M. & Bergles, D.E. Nature Neuroscience 16, 668–676 (2013).

20. Marisca, R. et al. Nature Neuroscience 23, 363–374 (2020).

21. Zhang, X. et al. Nat Commun 12, 5740 (2021).

22. Lam, M. et al. Nat Commun 13, 5583 (2022).

23. Pan, L. et al. Elife 12, e77441 (2023).

24. Culverwell, J. & Karlstrom, R.O. Semin. Cell Dev. Biol. 13, 497–506 (2002).

25. Pang, Z.P. & Südhof, T.C. Curr. Opin. Cell Biol. 22, 496–505 (2010).

26. Ros, O., et al. Cell Rep. 32, 107934 (2020).

27. Godoy, M.I., Pandey, V., Wohlschlegel, J.A. & Zhang, Y. bioRxiv 2024.07.22.604699 (2024).doi:10.1101/2024.07.22.604699

28. Frischknecht, R. & Gundelfinger, E.D. Adv. Exp. Med. Biol. 970, 153–171 (2012).

29. Sorg, B.A., et al. J. Neurosci. 36, 11459–11468 (2016).

## References

30. Vagionitis, S. et al. Cell Reports 38, 110366 (2022).

31. Freeman, J. et al. Nature Methods 11, 941–950 (2014).

32. Kwan, K.M. et al. Dev Dyn 236, 3088–3099 (2007).

33. Auer, F., Vagionitis, S. & Czopka, T. Current Biology 28, 549–559.e3 (2018).

34. Iyer, M., et al. Nat. Commun. 15, 265 (2024).

35. Distel, M., C, H.J. & Köster, R.W. Commun. Integr. Biol. 4, 336–9 (2011).

36. Coomer, C.E., et al. Nat. Neurosci. 1–12 (2024).doi:10.1038/s41593-024-01815-z

37. Meyer, M.P. & Smith, S.J. J. Neurosci. 26, 3604–3614 (2006).

38. Zhang, Y., Marshall-Phelps, K. & Almeida, R.G. Development 150, dev201323 (2023).

39. Sternberg, J.R. et al. Current biology: CB 26, 2319–2328 (2016).

40. Arganda-Carreras, I., et al. Lect. Notes Comput. Sci. 85–95 (2006).doi:10.1007/11889762_8

41. Dobin, A., et al. Bioinformatics (Oxford, England) 29, 15–21 (2013).

42. Gahtan, E., Tanger, P. & Baier, H. Journal of Neuroscience 25, 9294–9303 (2005).

43. Bene, F.D. et al. Science 330, 669–673 (2010).

44. Meyer, M.P. & Smith, S.J. Journal of Neuroscience 26, 3604–3614 (2006).

45. Portugues, R. & Engert, F. Frontiers in Systems Neuroscience 5, 72 (2011).

